# Testing the coordination hypothesis: incompatibilities in the absence of a single-cell bottleneck in an experimentally evolved social amoeba

**DOI:** 10.1101/2024.05.12.593719

**Authors:** Israt Jahan, Trey J. Scott, Joan E. Strassmann, David C. Queller

## Abstract

Multicellular organisms that form by aggregation of cells arguably do not achieve high levels of complexity. Conflict among the cells is a widely accepted explanation for this, but an alternative hypothesis is that mixing cells of different genotypes leads to failures of coordination, which we call the ‘coordination hypothesis’. We empirically tested the coordination hypothesis in the social amoeba *Dictyostelium discoideum*. We mixed *D. discoideum* clones that had evolved in isolation for generations and accumulated mutations that have not been tested against each other by selection. To quantify the effect of incompatibilities, we measured performance in terms of the developmental traits of slug migration and spore production. Importantly, kin recognition incompatibilities were avoided by mixing lines evolved from the same ancestor under conditions that would not select for the evolution of *de novo* recognition. Our results show no evidence of incompatibilities in coordinated movement of slugs towards light in the social amoeba. Spore production was higher than expected in mixtures, in apparent contradiction to the coordination hypothesis. However, we found support for coordination incompatibilities in an interaction between migration and spore production: in mixtures, fewer cells succeeded at migrating and becoming spores.

## Introduction

Many familiar life forms have evolved from groups of smaller, formerly free-living biological entities (Gardner and Grafen, 2009; Szathmáry and Maynard Smith, 1995; Bourke, 2013; Bell and Mooers, 1997; Corning and Szathmáry, 2015; Orr, 2000). The process of groups evolving a higher-level entity upon which natural selection can act is called a major transition. The evolution of multicellularity is one such major transition where single-celled ancestors evolved to take advantages of group living. Early multicellular life set the stage for a new type biological individual, allowing natural selection to operate at levels previously absent (Michod, 2005, 2007; Queller and Strassmann, 2009). How and why multicellular organisms developed and subsequently evolved, diversified, and increased in complexity, is an important question for understanding major transitions.

Multicellularity originated independently at least 20 times across all major domains of life (Grosberg and Strathmann, 2007). Eukaryotic multicellular organisms largely reproduce by undergoing single-cell bottlenecks. For example, sexually reproducing organisms produce gametes that combine into a unicellular zygote. Asexual plants and animals, such as the Amazon molly *Poecilia formosa* (Turner et al., 1980), several weevils (Suomalainen and Saura, 1973), and many angiosperms also reproduce by single-cell propagules (Bicknell and Koltunow, 2004).

In many organisms, alternative routes to multicellularity also exist where the offspring is likely to develop from more than one genotype. This type of reproduction may involve budding or fission(Galliot, 2012; Åkesson et al., 2001), or specialized structures, such as the fungal conidia or gemmae found in algae, mosses, and ferns (Hughes, 1971). Because the propagules are multicellular, they may include genetic variation that would be passed on to progeny. An extreme form of such chimeric multicellularity is perhaps best demonstrated by aggregation of individual cells that reside as neighbors (Jahan et al., 2022), known as aggregative multicellularity. The diversity of routes to simple multicellular development raises a fundamental question for the study of organismal complexity: Why haven’t other modes of multicellularity like aggregation equally contributed to complex development?

Single-cell bottlenecks are thought to be a requirement for complex multicellularity (Wolpert and Szathmáry, 2002; Fisher et al., 2020; Howe et al., 2024). The accepted explanation is kin selection (Hamilton, 1963; Axelrod and Hamilton, 1981; Bonner, 1988; Fisher et al., 2013). Development from a single-cell bottleneck results in a multicellular body with cells that are essentially genetically identical, minimizing genetic conflict and maximizing cooperation between cells within a clonal organism. When an exploiter mutant arises within the multicellular body during development, it might gain an initial advantage. But after a single-cell bottleneck, the mutant will exist in bodies with all mutant cells, and therefore unable to exploit further. A single-celled propagule thus reestablishes genetic uniformity for the germline, such that exploiters can exploit only in the lifespan of a single organism.

Wolpert and Szathmáry (2002) propose an alternative (but not mutually exclusive) explanation for why complex multicellular organisms develop from a single cell. They suggested that the development of complex multicellularity is not only a question of kin selection controlling conflict between cells, but also an issue of how cells coordinate developmental processes in the organism. For example, the authors suggest cells in chimeric mixtures could fail to generate repeatable developmental patterns:

> *“*… *patterning processes require signaling between and within cells, leading ultimately to gene activation or inactivation. Such a process can lead to reliable patterns of cell activities only if all the cells have the same set of genes and obey the same rules*.*”*

They add that this would hinder the evolution of novelty:

> *“*… *practically impossible to have several-to-many asexual, partly differentiated, cell lineages mutating in all sorts of directions in genetic space and yet keep up the ability to evolve into viable novel forms*.*”*

We call this idea the **“coordination hypothesis”**. Though Wolpert and Szathmary’s ideas may be more complex than our interpretation, we take the coordination hypothesis to include at least the following predictions. First, aggregative development puts new mutations into many new and evolutionarily untested cellular combinations. When this is so, we predict that mixing of cells from different lineages will usually be detrimental for the same reasons that untested mutations are generally detrimental. Second, even if a new advantageous mutation would be beneficial when fixed, to achieve fixation it must be advantageous across many frequencies and cell combinations. Even if some combinations were beneficial, evolution would be impeded if it had to pass through any stage with disadvantageous combinations. Specifically, we predict that this could occur when intermediate mixtures – groups where only some cells have a beneficial mutation – would have lower fitness than when it is present in all cells or none, that is when mixing produces transgressively lower fitness. In contrast, organisms with a single-cell bottleneck do not face these problems because each organism is genetically uniform and the mutation will fix if it is beneficial when present in all cells in an organism.

To make this logic more concrete, consider how evolution might work on human bodies if they were formed by aggregation of diverse cells. A new mutation might find itself first in the liver, but in the next generation it might find itself in the hypothalamus of one individual, the adenoids of another, and the germ cells of a third. To rise to higher frequencies, it would have to be advantageous, on average, across all these contexts. If the frequency did increase, it would find itself in still more localities and combinations of localities. The complexity increases much more if there are other segregating mutations.

In Figure 1, we present a schematic diagram of possible outcomes when cells from distinct lineages are mixed together. The null hypothesis (Figure 1, gray dotted line) predicts that mixing is neither a beneficial nor detrimental to multicellularity, such that the fitness of the mixture is the average of the fitness of the two lineages A and B (shown by the gray circle in both panels). This would not impede evolution from the low-fitness B to the high-fitness A. The coordination hypothesis predicts that mixing will have detrimental fitness consequences in the resulting multicellular group (Figure 1, line 1) on average compared to the null expectation. In some extreme cases (Figure 1, line 2), incompatibilities between cells may result in large fitness costs for the mixture causing it to have transgressively lowest fitness. This would imply that a clone starting with a lower fitness (as is the case for Clone B) would not be able to evolve to the higher-fitness A state because it would require passing through a lower fitness stage. Under this scenario, intermediate cell combinations would hinder the evolution of increased complexity, as Wolpert and Szathmary argued.

**Figure 1.**
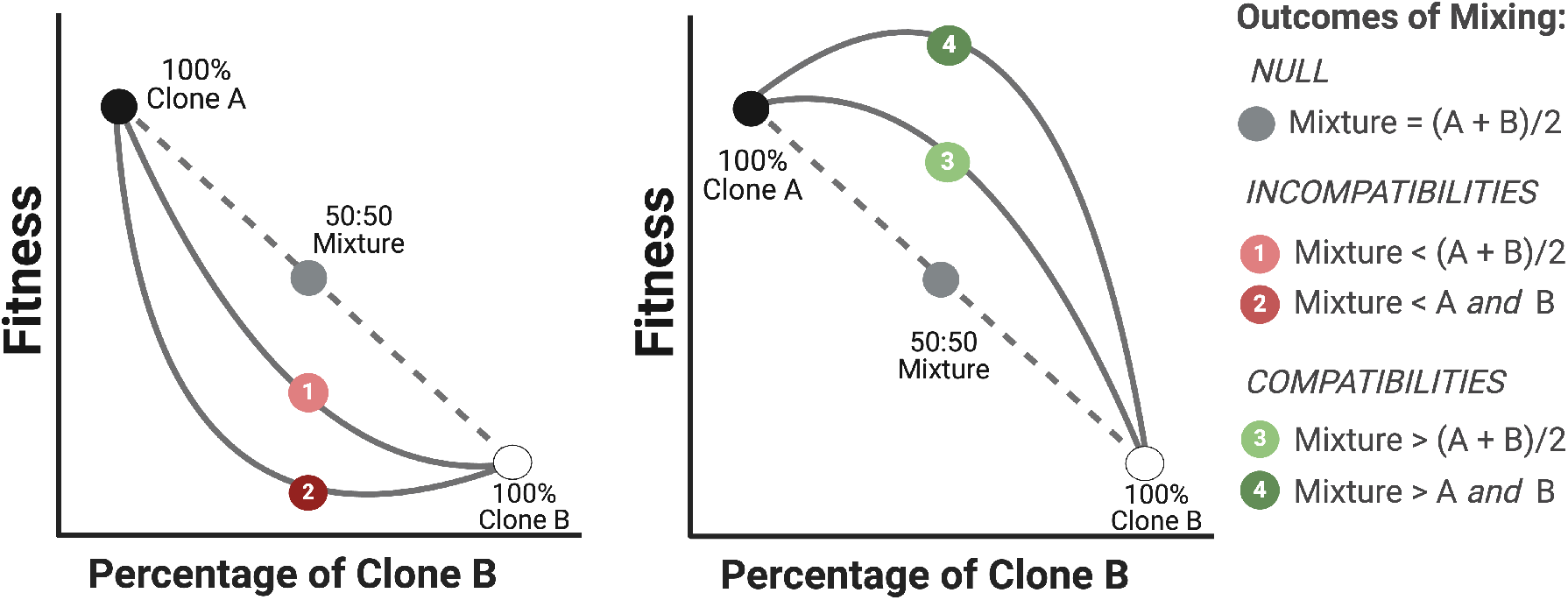
Illustration showing the possible fitness outcomes of mixing between two cell lineages. The gray dotted line represents the Null hypothesis (additivity). The panel on the left shows the two predictions of coordination hypothesis, where incompatibilities in development may impede fitness improvement. The panel on the right shows the possibility of mutations that are compatible and may generate beneficial fitness consequences upon mixing. Circles represent the 50:50 mixtures that will be tested in this study

Alternatively, mixing could sometimes result in improved fitness in the mixtures compared to the expected null (Figure 1, line 3 and 4). Under some conditions, for example social heterosis (Nonacs and Kapheim, 2007), mixing may generate combinations that are complementary in nature such that each lineage compensates for genetic defects in the other and results in higher or even transgressively higher fitness in the mixture. Though social heterosis is often defined for mixtures of multicellular individuals, such as ants, such complementarity can also apply to microbes (Kraemer and Velicer 2014). Under social heterosis, mixing is predicted to be favored by natural selection, other things being equal (Nonacs, 2017; Ebrahimi and Nonacs, 2021).

To test the coordination hypothesis, we require a biological system that can be modified to 1) develop from mixtures of genetically different cells (chimeric development), 2) provide developmental mutations for mixing that have not been previously tested by selection, and 3) exclude kin recognition as a cause of differences in mixtures. The social amoeba *Dictyostelium discoideum* can satisfy these criteria and therefore is an ideal system for this experiment.

Individual *D. discoideum* amoebae divide and grow independently in the presence of abundant edible bacteria but under starvation they aggregate and undergo multicellular development (Figure 2). At one stage in the development, the multicellular aggregate forms a slug that shows coordinated movement towards light. Migrating towards light can assist *D. discoideum* in moving towards the soil surface (Castillo et al., 2005) and likely increases the chance of dispersal by insects, other small invertebrates, and birds (Huss, 1989; smith et al., 2014). The slug culminates into a fruiting body with some cells becoming spores held atop a slender stalk made up of dead cells. Because of this lifecycle, *D. discoideum* has been extensively used to study chimeric development (Mehdiabadi et al., 2006; Khare and Shaulsky, 2006; Khare et al., 2009; Sathe et al., 2010; Castillo et al., 2011; Foster, 2010).

**Figure 2.**
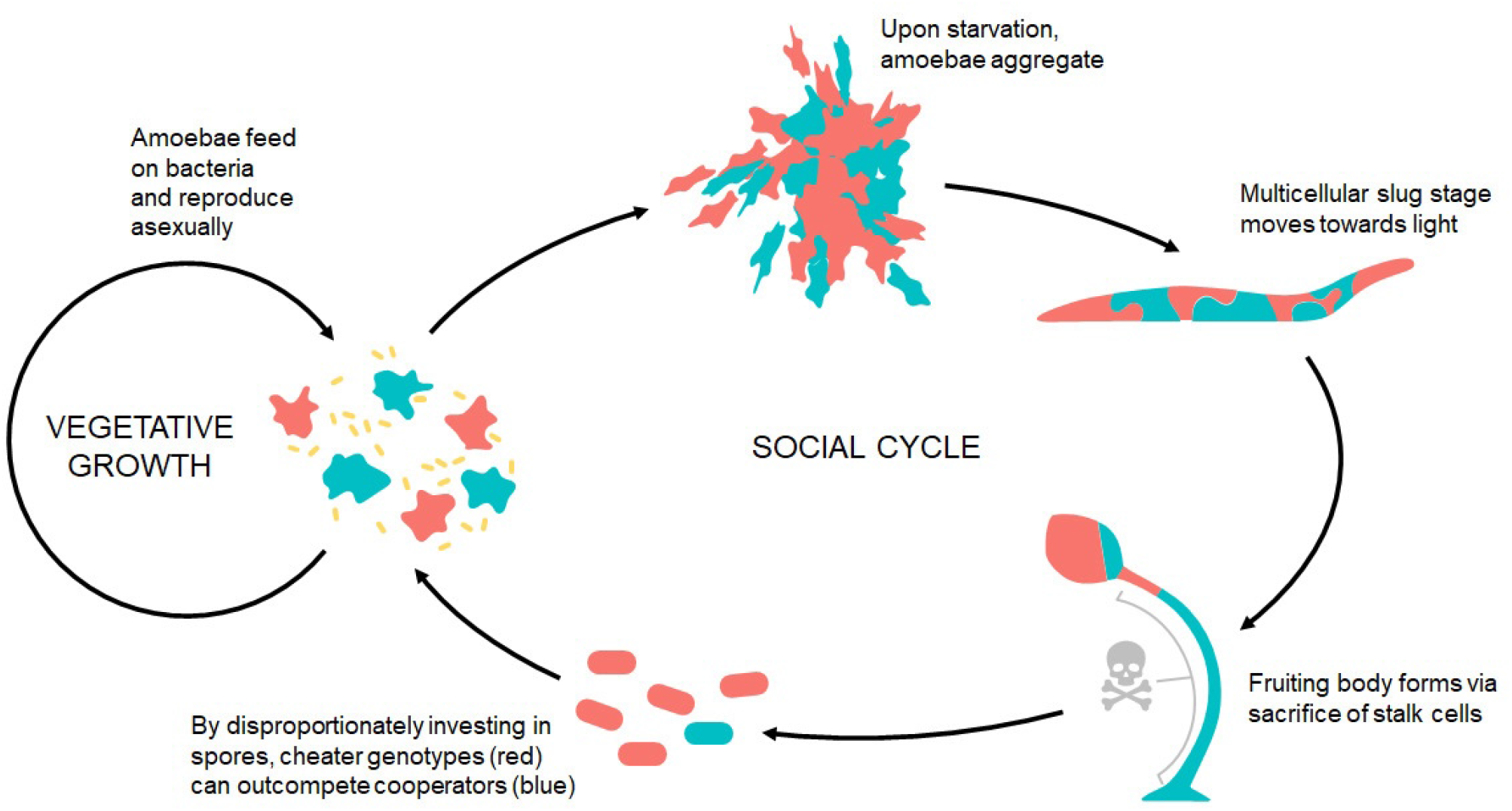
Simplified life cycle of *D. discoideum* illustrating potential for cheating. Credits Tyler Larsen, CC BY-SA 4.0, via Wikimedia Commons.

There are some documented costs of mixing with non-relatives (Strassmann et al., 2000; Foster et al., 2002; Fortunato et al., 2003). Genotypes in chimeras could invest less in dead stalk production while maximizing the fitness benefits of producing more spores (Santorelli et al., 2008; Khare and Shaulsky, 2010; Buttery et al., 2010; Noh et al., 2018). Chimeric slugs move shorter distances than clonal slugs when started with the same number of cells (Foster et al., 2002). This could result from competition among the genotypes to avoid the pre-stalk region located in the front of the slug by moving towards the posterior pre-spore region. Most known cases of costs of chimeric development in *Dictyostelium discoideum* have been studied by mixing natural strains that that have been under selection over a long evolutionary timescale, and arguably could have evolved a kin recognition system.

To test for unselected incompatibilities, we need to exclude these evolved strategies and any underlying kin recognition. At least in the early stages of aggregation, a matching pair of cell surface receptors encoded by the *tgrB1* and *tgrC1* alleles is necessary and sufficient for attractive self-recognition in *D. discoideum* (Hirose et al., 2011; Ho et al., 2013; Hirose et al., 2017). Amoebas can selectively bind to other cells with compatible *tgr* alleles such that a mismatch can cause poor cell-cell adhesion, and even assorting into individual clones later on (Ostrowski et al., 2008; Benabentos et al., 2009; Hirose et al., 2011; Ho et al., 2013; Hirose et al., 2017), although sorting may largely disappear at later stages, perhaps due to slug fusion.

Eliminating kin recognition can be achieved by using clones evolved from a common ancestor without the multicellular stage. In a recent experimental evolution study (Larsen et al., 2023), we created such lineages of *Dictyostelium discoideum* clones. Amoebas were transferred on nutrient plates before the onset of starvation, for 30 transfers (more than 200-300 cell divisions). With abundant nutrients, these *D. discoideum* lines did not form multicellular groups and would therefore have experienced no selection for novel kin recognition variants. However, these lines did evolve altered social behaviors via drift or pleiotropy with traits under selection: they showed reduced cheating, slug migration, and increased spore production. In sum, we have experimental lines that should have the same kin discrimination genotypes but differ genetically in social traits that have never been selectively tested in combination. Here we use these lines to generate chimeric mixtures without kin recognition to test for a reduced coordination in slug migration and spore production.

## Methods

### Chimeric Mixes

Our study involved two kinds of mixtures: pairwise and complex. First, we made pairwise mixes of clonal ancestors with one of their evolved lineages. We used 10 *D. discoideum* ancestors (QS6, QS9, QS11, QS18, QS69, QS70, QS159, QS161, QS395, QS859) that had been experimentally evolved in triplicate to generate 30 evolved lines in total (Larsen et al., 2023). We made pairwise mixes of the ancestral clones (A) with each of their own derived lines (E1, E2, E3) to generate a total of 30 mixed lines (A+E1, A+E2, A+E3; we call these “lines” for consistency, but they are really mixtures). Therefore, we had a total of 70 experimental lines from 10 *D. discoideum* clones. We had 3 technical replicate plates for each of the 70 experimental lines resulting in a total of 210 plates for 3 Treatments – Ancestor, Evolved, and Mixed.

For generating a complex mixture, we mixed the three evolved lineages derived from a common ancestor with each other for 3 out of 10 strains (QS6, QS9, QS18). There should be twice as many differences between two evolved lines as between ancestor and evolved, generating greater power in detecting incompatibilities, should they occur. Mixed lines from complex mixtures did not include the ancestors. We repeated the experiment with complex mixtures on 3 different days to enhance power, resulting in a total of 90 plates for the analyses (30 plates X 3 days). On each day, we had two replicate plates for every ‘Unmixed’ evolved line (E1 or E2 or E3) and four replicate plates for every ‘Mixed’ treatment (equal proportions of E1 and E2 and E3) resulting in a total of 10 plates for every strain.

### Slug Migration Assay

To obtain spores to inoculate the slug migration assays, we first took glycerol stocks of *D. discoideum* spores stored at 80 °C and streaked each frozen sample on solid SM/5 agar plates (Fey et al., 2007). After development, we collected spores and plated 2 × 10^5^ of them onto a fresh SM/5 plate along with 200μl *K. pneumoniae* food bacteria resuspended in KK2 buffer (OD600=1.5). This round of growth was to remove freezer effects. On day 4 we harvested spores from developed fruiting bodies, and initiated the slug migration assay on non-nutrient agar plates (13 cm diameter), in three replicates for each ancestral clone, their evolved lineages, and mixtures.

On every plate, we marked a 10 cm secant line (henceforth inoculation line) on which we loaded spore samples (Figure 3). For pairwise mixtures, we loaded sample consisting of 10^7^ spores in 50μL of *K. pneumoniae* (OD600=50.0) suspended in KK2 buffer. The ‘Mixed’ treatment for pairwise mixtures were made of 50% of Evolved (E1 or E2 or E3) and 50% Ancestor (A) spores in the total 10^7^ spore suspension. For samples of complex mixtures, we loaded 9 × 10^6^ spores suspended in 50 μL of *K. pneumoniae* food bacteria (OD600=50.0) on the inoculation line as in pairwise mixtures. We repeated the experiment on 3 different days and collected data from a total 90 plates.

**Figure 3.**
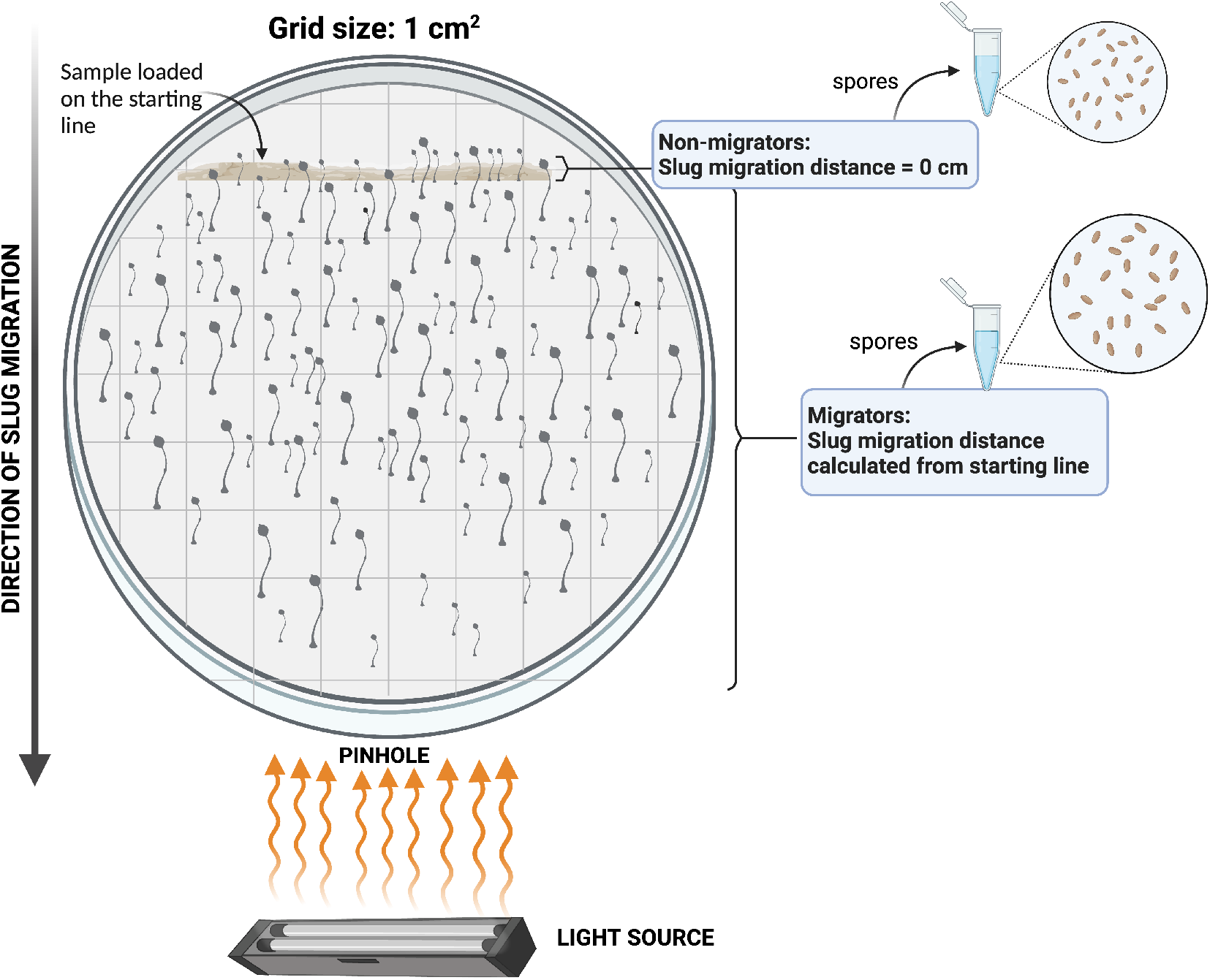
Schematic depiction of an experimental plate at the end of the slug migration assay. We refer to fruiting bodies that developed directly on the inoculation site as non-migrators and fruiting bodies that developed beyond the inoculation line as migrators. Fruiting bodies on a plate, especially non-migrators, were more numerous than shown in the diagram. Distance travelled by migrators towards light source was measured from the inoculation line. We counted spores from both migrator and non-migrator fruiting bodies and added the two to obtain total spores produced on a plate. Image created with BioRender. Objects not drawn to scale.

After loading the suspension along the starting line of the agar plates, we allowed the sample to dry and wrapped the plates individually in aluminum foil. We made a small pinhole opposite the starting line on each wrapped plate. For the duration of the experiment, light enters only through this small pinhole towards which slugs migrate from the other end of the plate. We then left the wrapped plates undisturbed for 8 days with the pinhole side of the plate facing a light source in the laboratory. Because the plates had no nutrients, bacteria and amoebas grew only at the starting line, but slugs could migrate off the starting line towards the light. We unwrapped the plates at the end of the 8 days, and allowed fruiting bodies to finish developing fully under direct light.

### Image Analysis

We used a Canon EOS 5D Mark III camera to photograph each plate. We put each plate on a laboratory bench with the camera mounted at a fixed distance. We obtained slug migration distances using the software Fiji and ImageJ (Bourne and Bourne, 2010; Schindelin et al., 2012; Rueden et al., 2017). We processed each image by first scaling it and then overlaying a 1cm × 1cm grid. We recorded the distance of each fruiting body from the starting line. For pairwise mixes, we assigned distance=0 cm for fruiting bodies that developed directly on the inoculation line, and refer to those as ‘non-migrators’. Fruiting bodies that traveled beyond the inoculation site are referred to as ‘migrators’ (as shown in Figure 3).

We observed a few fruiting bodies traveled in the opposite direction to light. To account for such slugs that may contribute to a reduced average migration distance of slugs on a plate, we measured the distances of all fruiting bodies from the inoculation line on a plate in the follow-up experiment. Our measurements for complex mixtures therefore account for positive or negative movement towards light for slugs on a plate.

### Spore Production Assay

At the end of the slug migration assay for pairwise mixtures, we quantified the number of spores produced on a plate by slugs that migrated towards light as well as those that did not leave the inoculation site. We collected non-migrators and migrators for each plate in separate eppendorf tubes (Figure 3) containing 1mL KK2 buffer. We made 1:100 dilutions of the collected spores and counted them using a hemacytometer. We added the number of spores produced by migrators and non-migrators to obtain the total spore production on a plate for each strain across three treatments. We performed a log transformation for use in our analyses. This assay was performed only for pairwise mixtures and not for complex mixtures.

### Statistical Analysis

We used R version 4.2.1 (R Core Team, 2022) for all our analyses. We used the *tidyverse* (v2.0.0) suite of packages for data cleaning (Wickham et al., 2019), and the package *fitdistrplus* (v1.1.11) for fitting univariate distributions to our data (Delignette-Muller and Dutang, 2015). For both pairwise mixtures and complex mixtures, we performed linear mixed effects modelling with the *lme4* (v1.1.35.2) package (Bates et al., 2015).

Treatment and Clone were fixed effects, and replicate line a random effect for the analysis of both slug migration and spore production data (see Table 1 and Table 2 for details on experimental lines). Pairwise mixtures included three treatments (Ancestor, Mixed, and Evolved) whereas complex mixtures included two treatments (Mixed and Unmixed). We used Akaike Information Criteria (AIC) for model selection which estimates the relative quality of each model based on model quality and parsimony. To assess model fit and assumptions, we used the package *performance* (v0.11.0) (Lüdecke et al., 2021).

We fitted a generalized linear mixed effects model to our proportion data from pairwise mixtures using the package *glmmTMB* (v1.1.9) with a beta-binomial distribution (Brooks et al., 2017). For the analysis of slug migration distances, we used the *robustlmm* (v3.3.1) package (Koller, 2016) to account for outliers without their removal from the data used in our final model. A robust analysis of our final mixed effects model prevents outliers from influencing estimations by inversely weighing them. We reported estimated marginal means averaged at the level of clone using the *emmeans* package (v1.10.1) (Lenth, 2022) for all the statistical models. We performed hypothesis testing using the estimated marginal means and inferred statistical significance of our results after correcting for false discovery rates. All graphs in the results section were plotted with the *ggplot2* (v3.5.1) package (Wickham, 2016) and significance values were layered using the *ggsignif* (v0.6.4) package (Constantin and Patil, 2021).

## Results

### Slug migration in mixtures is not different from the mean of the unmixed populations

We quantified the outcome of mixing on slug migration in 10 strains of *D. discoideum*. First, we created pairwise mixtures between clonal ancestors and their experimentally evolved descendants for the three treatments: Ancestors, Evolved, and Mixed (50% Ancestor + 50% Evolved). We measured average slug migration distance for all fruiting bodies on a plate (see Figure 3 for migrators and non-migrators). For pairwise mixtures, chimeras were not significantly different from the expected value on average ([Ancestor + Evolved]/2 – M = 0.172, SE= 0.113, z.ratio = 0.1528, p.value = 0.19), as shown in Figure 4).

**Figure 4.**
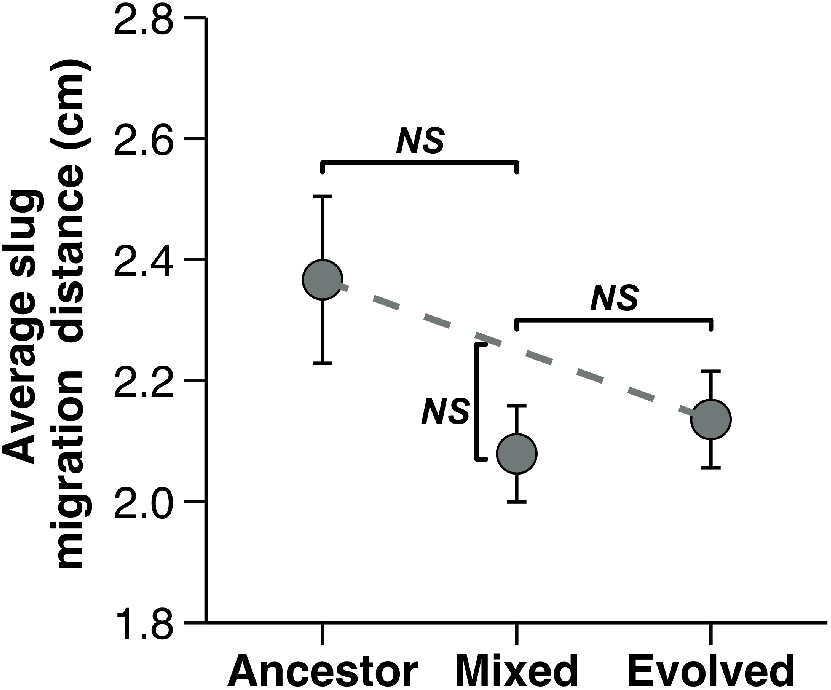
Average slug migration distance on a plate across three treatments. The expected averages of pairwise mixtures between ancestor and evolved lines are depicted with a dashed line, against which the observed mixed point is compared. Error bars correspond to standard error of the estimated marginal mean. Slug migration in mixtures, on average, was not significantly different from the Ancestors, the Evolved lineages, or from the expected value.

The average slug migration distances depicted in Figure 4) are composed of two components, the fraction of slugs that migrated and the distance traveled by those that migrated (as shown in Figure 3). We analyzed these separately. First, there was no significant difference between the expected and the observed values of proportion of migrators on a plate ([Ancestor + Evolved]/2 – M = 0.01735, SE= 0.0169, z.ratio = 1.026, p.value = 0.4571 (Figure 5), panel A). We then compared slug migration distances for only the migrators for the three treatments (Figure 5), panel B). Mixed lines were extremely close to the expected value ([Ancestor + Evolved]/2 – M = -0.00346). A post hoc comparison using a planned emmeans contrast, averaged over the level of clones, showed no significant difference (SE = 0.16, z.ratio = -0.022, p = 0.9828).

**Figure 5.**
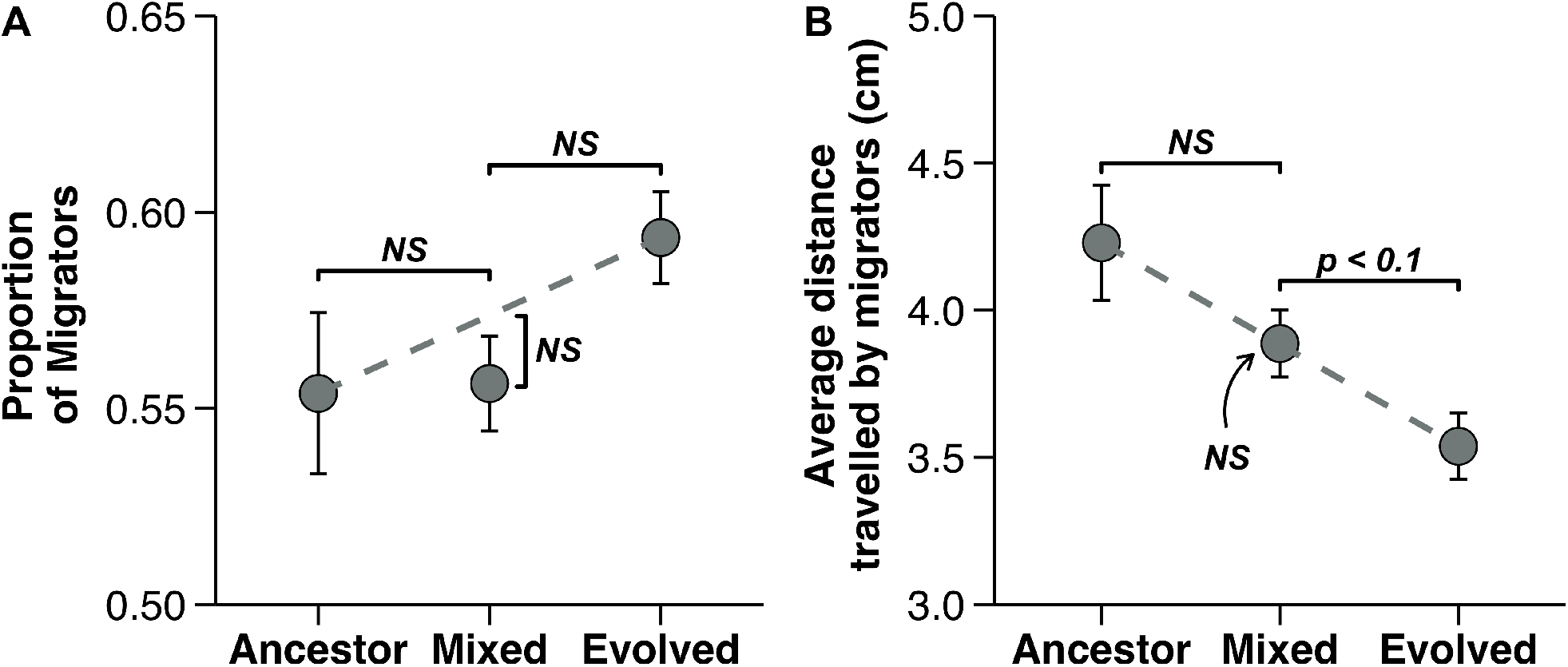
Components of slug migration (A) Proportion of migrators across three treatments averaged at the level of clones. There is no significant difference in the proportion of migrators on a plate among the treatments. (B) Estimated average slug migration distance of migrators on a plate. Pairwise mixtures travel an intermediate distance that is not significantly different from the expected value. Mixtures, however, travel a greater distances than the evolved lineages. Mixed fruiting bodies that migrated beyond the inoculation line were not significantly fewer than the expected average of Ancestors and Evolved lines.

We performed a second slug migration experiment with the aim of getting increased power to detect any incompatibilities. We mixed three descendants (from the same ancestor) to obtain a more complex mixture that should allow greater statistical power and thus robust inference. For two reasons, we expect the mixture of three evolved lines to include more genetic differences than the pairwise mixes. First, more cell variants are mixed together. Second, two descendants have twice as many differences as ancestor and descendant. Our analysis still showed no significant differences in slug migration distances between unmixed and mixed evolved lineages of a clonal ancestor (Unmixed – Mixed = -0.00976, se =0.197, z. ratio = -0.050, p. value = 0.9604) as shown in Figure 6.

**Figure 6.**
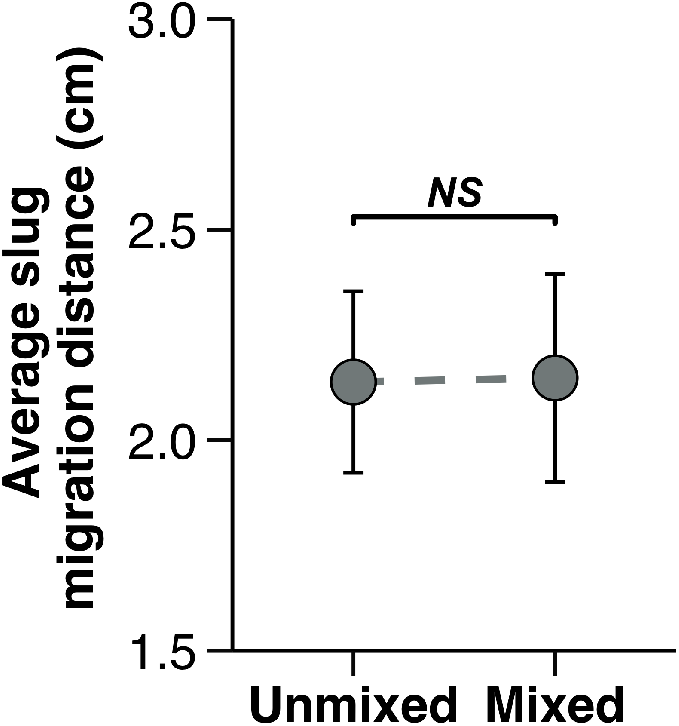
Complex mixes of evolved lineages of a clonal ancestor are not statistically different from evolved lines in isolation for average slug migration distance. Data are averaged at the level of clones (QS6, QS9, QS18).

### Spore production in pairwise mixtures is higher than the expected average of ancestor and evolved lines

We measured the number of spores collectively produced by developed fruiting bodies from pairwise mixes at the end of slug migration assay. We found that mixed lines, on average, produced significantly more total spores compared to the expected average of ancestral and evolved lines of *D. discoideum* ([Ancestor + Evolved]/2 – M = -0.1563, SE = 0.0477, z.ratio 7. Moreover, the mixed lines produced significantly higher number of spores than the ancestral lines (Ancestor – Mixed = -0.2474, SE = 0.0675, z.ratio = -3.664, p.value = 0.0007).

### Migrating slugs produced transgressively lower spores in mixtures

We further assessed a kind of interaction between slug migration and spore production by quantifying the proportion of spores produced by migrators with respect to total spore production on a plate. At the end of migration, cells in slugs can get into the next generation by successfully producing spores. The proportion of spores produced by migrators was significantly less than the expected average of ancestors and mixed lines ([Ancestor + Evolved]/2 – M = 0.0413, SE = 0.0109, z.ratio = 3.790, p.value = 0.0005), as shown in Figure 8.

Mixtures produced significantly lower number of spores than both the ancestors (Ancestor – Mixed = 0.0461, SE = 0.0158, z.ratio = 2.925, p.value = 0.0034), as well as the evolved lineages (Evolved – Mixed = 0.0366, SE = 0.0108, z.ratio = 3.386, p.value = 0.0011).

Though migrating slugs contributed fewer spores to the total number of spores produced on a plate across treatments, mixtures appeared to incur the most cost of migration.

## Discussion

Complex multicellular life more commonly arises in lineages with single-celled bottlenecks (Grosberg and Strathmann, 2007; Fisher et al., 2020; Howe et al., 2024). This may be because of reduced conflict when the multicellular body consists of close relatives. An alternative explanation, which call the coordination hypothesis, is that mixing of genetically distinct cells with different mutations in novel, untried combinations, is expected to have a detrimental effect on the developing multicellular organism due to incompatibilities in coordinating development. The presence of incompatibilities could help explain why aggregative organisms are relatively rare and less complex compared to clonally developing multicellular organisms (Wolpert and Szathmáry, 2002).

We investigated whether mixing experimentally evolved lineages (with ancestors, and among themselves) with mutations that have not been tested against each other by selection would result in detrimental effects in the multicellular stage of an aggregative social amoeba. We predicted detrimental consequences of mixing could manifest in two different ways. First, mixing between two isolated lineages can lower fitness on average. Second, mixtures could have lower fitness than both constituent cell lineages such that mixtures have transgressively lower fitness. We looked for such developmental incompatibilities in *Dictyostelium discoideum* by measuring slug migration distance and spore production.

With respect to slug migration distance, we observed no evidence for incompatibility. There was no significant differences across treatments for average slug migration distance on a plate (Figure 4). To test whether non-migrators were disproportionately influencing migration distance across treatments, we compared the treatments for distance travelled by the migrators only. We found no difference in the proportion of slugs migrating towards light across treatments in pairwise mixtures (Figure 5A). We further observed that average distance travelled by migrators in pairwise mixtures was nearly exactly intermediate between the ancestors and their evolved descendants (Figure 5B). To increase power of detecting incompatibilities, we made complex mixtures of three descendant lines of a clonal ancestor. These complex mixtures also did not show any evidence for incompatibilities (Figure 6). These results fail to provide any support for the hypothesis that previously unselected combinations of genotypes would be deleterious.

Spore production presents a different picture. The spore number of mixtures was significantly higher than the average of pairwise mixtures and the ancestors, but was not significantly higher than the evolved lineages (Figure 7). The increase in total number of spores produced in mixtures from the expected value may appear to be opposite to the incompatibility hypothesis, if we assume that higher spore production results in higher fitness. Spores are often used as a measure of fecundity (Kuzdzal-Fick et al., 2023; Scott et al., 2022; Buttery et al., 2009), and therefore our spore production result would appear to represent an improvement in fitness upon mixing. However, an increase in spore production may not necessarily result in improved fitness if it is generated at the cost of stalk formation or by producing smaller spores. Support for this possibility comes from the experiment that generated these lines (Larsen et al., 2023). Because there was no social stage during the experimental evolution, we expected most aspects of social fitness to change negatively, and both the ability to cheat and the ability to migrate did decline as expected. However, spore production increased, suggesting, but certainly not proving, that higher spores production represented a decline in fitness.

**Figure 7.**
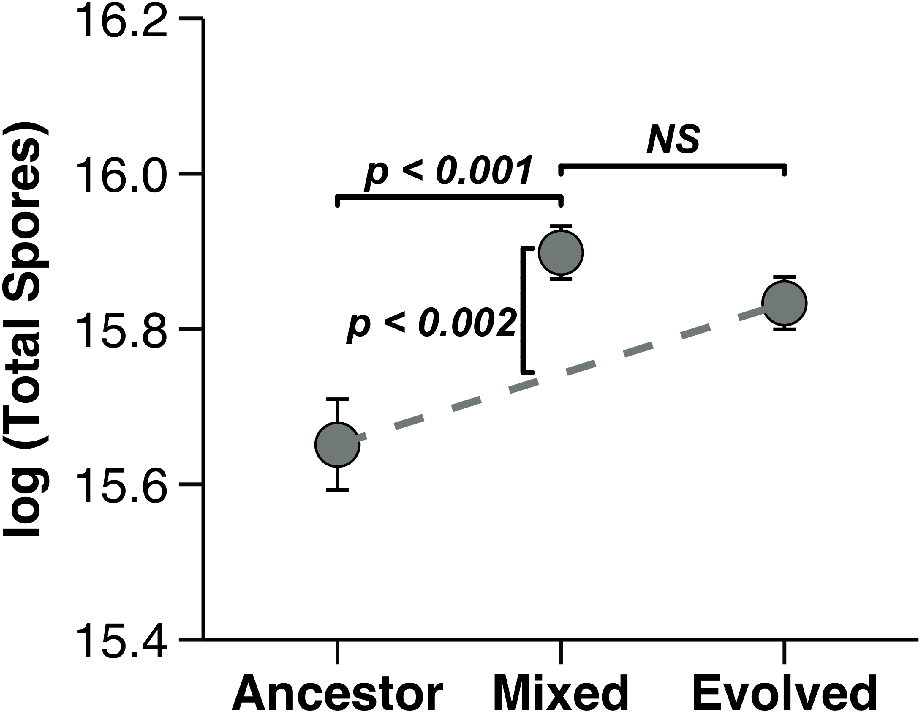
The log transformed estimate of total spores produced by mixed lines on average is significantly greater than the expected average when ancestors and their evolved counterparts are mixed in equal proportions in *D. discoideum*.

**Figure 8.**
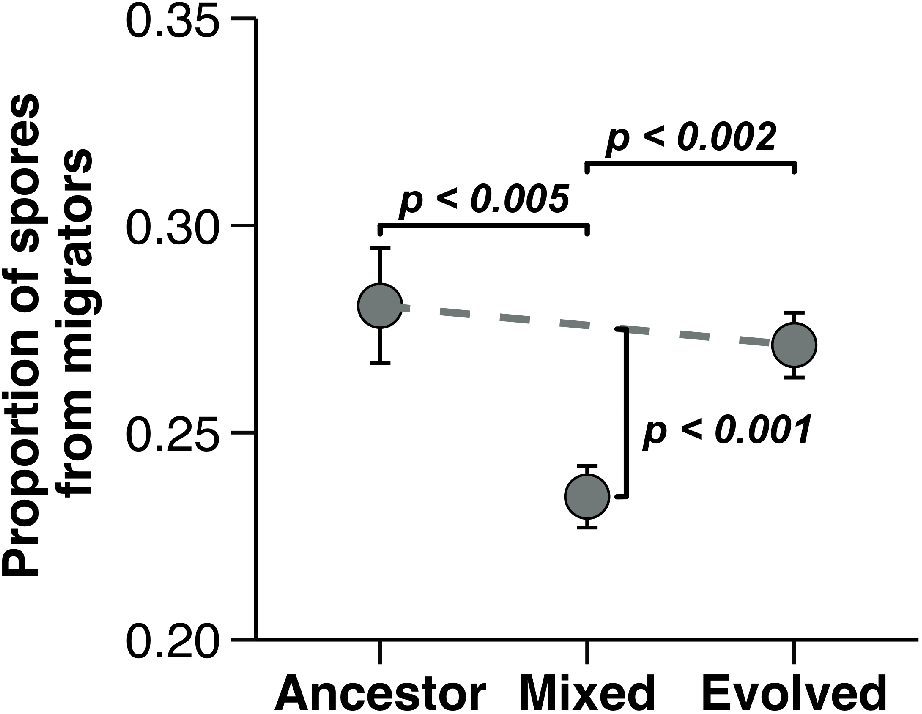
Slugs that migrated towards light in mixed lines on average show significantly lower contribution towards total spores produced than both ancestors and their evolved counterparts. p-values shown in the graph were obtained by Post hoc pairwise comparisons of the ratio of migrator spores to total spores produced on a plate. We used Z-tests, corrected for false discovery rates.

More convincing evidence for a transgressive decline in fitness comes from a sort of interaction between migration and spore number. We could not look at the interaction at the individual slug level because we did not count spore numbers for individual migrating slugs. However, we collected and counted total numbers of spores for migrating and non-migrating slugs from pairwise mixtures. Mixtures showed transgressively lower proportions of total spores coming from migrating slugs (Figure 8) with respect to total spores produced on a plate. Thus, we saw no significant incompatibility in the proportion of slugs migrating (Figure 4), there is incompatibility when the migrating proportion is expressed in terms of ultimate spore number. Since phototactic migration improves fitness, mixing is extremely detrimental because it results in a smaller fraction of spores gaining that advantage. If transgression in spore production is disadvantageous, as we argue here, then our result supports the argument that incompatibilities in mixtures could prevent evolution from the lower fitness state to the higher fitness one.

In our study, there should have been no kin discrimination since the mixed lines were derived from the same clone. Moreover, social stages did not occur in the evolving lines (Larsen et al., 2023),so there was no selection for kin recognition for avoiding variants who could be cheaters. As expected in the absence of selection for cheating, cheating ability declined (Larsen et al., 2023). The relatively short duration of our experimental evolution, the large population sizes, and a very low mutation rate in *D. discoideum* (Kucukyildirim et al., 2020) make it very unlikely that genetic drift could cause changes in the *tgr*B1 and *tgr*C1 recognition loci across multiple lines. We are therefore confident that our results are not driven by the evolution of kin recognition loci.

Data from other studies of chimeric mixtures do not address the coordination hypothesis as clearly because they are likely complicated by kin recognition. Previous studies in *D. discoideum* have created chimeras by mixing naturally isolated wild genotypes with each other, and compared them to clonal isolates (Foster et al., 2002; Castillo et al., 2005; Jack et al., 2011, 2015; Kuzdzal-Fick et al., 2023). For example, Foster et al. (2002) showed that slug migration distances are lower in chimera compared to clonal slugs. This would appear to support the coordination hypothesis. But it could be the tgr recognition system, which operates via adhesion cells of the same genotype (Ostrowski et al., 2008, 2015; Benabentos et al., 2009; Hirose et al., 2011; Ho et al., 2013; Hirose et al., 2017).

A study quite similar to ours used the social bacterium *Myxococcus xanthus*, which cooperatively forms multicellular structures. Lines clonally evolved from a common ancestor showed incompatibilities in merging and swarming together when they were mixed (Rendueles et al., 2015). The authors favored the interpretation that these incompatibilities were not directly selected, which would make the results consistent with the coordination hypothesis. However, the evolving *M. xanthus* lines were allowed to socially swarm and form fruiting bodies so one cannot rule out direct selection for cheater resistance.

Rendueles et al. also suggested that, if the incompatibilities were not selected by kin recognition, then they would be very similar to Bateson-Dobzhansky-Muller incompatibilities, which cause isolated populations to be increasingly sexually incompatible and eventually become separate species (Orr, 1996; Dobzhansky, 1934; Bateson, 2009; Turelli and Orr; Turelli et al.). As one population evolves through natural selection, newly emerged genes are tested against existing genes in the population and selected to be compatible with the background. Novel genes that cause mismatches within a population are weeded out, but new genes in one population are never tested with genes that may have newly emerged in another isolated population. Therefore, mismatches between populations gradually accumulate and only become visible to natural selection when the populations later come into contact. For the coordination hypothesis, the incompatibilities must occur between interacting cells, arise within one population, and emerge on a much shorter time scale.

The prevalence of a single-cell bottleneck in multicellular lineages, particularly in complex multicellular organisms, could be due to the importance of high relatedness to cooperation among cells (cooperation) or to the importance of genetic uniformity for coordination. There seems little doubt that kin-selected cooperation is important () (Katoh-Kurasawa, Lehmann, and Shaulsky 2024; Kay, Keller, and Lehmann 2020; West et al. 2021), but it has been much harder to test the coordination hypothesis, particularly because evolved recognition effects are hard to exclude. We excluded them by mixing clonal lines evolved from a single ancestor under non-social conditions that should prevent selection for kin recognition. Our results provide some support for the idea that mixture of cells may be poor at coordinating with each other, and the transgressive effects support the idea that evolution may be impeded.

## AUTHOR CONTRIBUTIONS

I. J., D. C. Q. and J. E. S. conceived of the study. I. J. collected experimental data, and performed the statistical analyses with inputs from co-authors. All authors participated in writing and editing the manuscript.

## ACKNOWLEDGMENTS

We are thankful to the members of the Queller-Strassmann Lab for their feedback on this study. We thank Tyler. J. Larsen for the *D. discoideum* experimental evolution lines and insightful comments on the manuscript.

## DATA AVAILABILITY

Code to reproduce the results and plots in this study are in a Github repository (www.github.com/jahanisrat/WolpertHypothesis). Data used for statistical analyses is available in the in the submitted repository. Raw images used for this study are archived on Zenodo (10.5281/zenodo.12460200).

